# 222-nm far UVC exposure results in DNA damage and transcriptional changes to mammalian cells

**DOI:** 10.1101/2022.02.22.481471

**Authors:** Qunxiang Ong, Winson Wee, Joshua Dela Cruz, J. W. Ronnie Teo, Weiping Han

## Abstract

Ultraviolet (UV) germicidal tools have recently gained attention as a disinfection strategy against the COVID-19 pandemic but the safety profile arising from their exposure have been controversial and impeded larger scale implementation. We compare the emerging 222-nm far UVC and 277-nm UVC LED disinfection modules with the traditional UVC mercury lamp emitting at 254 nm to understand their effects on human retinal cell line ARPE-19 and HEK-A keratinocytes. Cells illuminated with 222-nm far UVC survived while those treated with 254-nm and 277-nm wavelengths underwent apoptosis via JNK/ATF2 pathway. However, cells exposed to 222-nm far UVC presented the highest degree of DNA damage as evidenced by yH2AX staining. Globally, these cells presented transcriptional changes in cell cycle and senescence pathways. Thus, the introduction of 222-nm far UVC lamps for disinfection purposes should be carefully considered and designed with the inherent dangers involved.

## Introduction

It is well established that mammalian cells can sense ultraviolet (UV) irradiation and mount a series of elaborate responses, also known as the UV response, which targets multiple signalling pathways^1^. These could eventually lead to apoptosis and local DNA damages^2^. The long-term negative consequences of the UV irradiation include erythema^3^, skin cancer^4^ and eye damages^5^.

Traditionally, we have focussed our attention on UVA (330 - 400 nm) and UVB (290 - 330 nm) given that these are the UV wavelengths we are typically exposed to on the earth’s surface. UVC (200-280 nm) is believed to be typically absorbed by the atmospheric ozone and does not reach the earth’s surface^6^. However, with increasing use of man-made UV germicidal irradiation tools, it is likely that more human exposures of UVC could occur. This is especially relevant during the COVID-19 pandemic, where UVC light manufacturers have introduced several UVC devices intended for home usage and safety precautions have been hugely neglected in this process.

Various studies have documented the potential dangers of UVC mercury lamp (254 nm), where temporary eye and skin damage could be observed upon prolonged direct exposure to UVC^7^. Irradiation of mammalian cells with UVC has been known to induce DNA damages in the form of thymine dimers and 6,4-photoproducts^1,8^, activation of MAPK signalling pathways^9,10^, and potentially cell cycle arrest and apoptotic pathways^2^. Recent developments in the UV technologies have seen UVC LED, mainly in the wavelengths of ∼270-280 nm^11^, and 222-nm far UVC lamp^12,13^ emerging as viable alternatives to the traditional UVC mercury lamp. However, their safety profile on mammalian cells have been much less documented, with that of 222-nm wavelength being controversial. Promisingly, 222-nm far UVC lamp has been touted to be a safe solution, where no pyrimidine dimer formation was observed in human and mouse skin models, and cell viability of layered cell sheets is retained after irradiation with 222-nm far UVC ^12^. However, some reports emerge that 222-nm sources may not be as safe where it is capable of inducing both erythema and cyclopyrimidine dimer (CPD) formation in human skin^14^.

Here, we sought to compare the different UVC wavelengths in their effects on cell viability and DNA damage. We found that human retinal cells, ARPE-19 cells, have decreased viability and lower growth rate when exposed to 222-nm far UVC compared to control cells. They exhibited cell viability with the JNK/ATF2 pathway remaining suppressed compared to 254-nm and 277-nm treatment. However, they had the highest DNA damage as evidenced by yH2AX staining. RNA sequencing of cells exposed to 222-nm far UVC showed that the cell cycle machinery is being disrupted, with DNA damage pathways and senescence pathways activated.

## Results

### UVC-induced cell viability reduction and apoptotic processes

We first examined the viability of ARPE19 cells exposed to different UVC wavelengths *via* the sulforhodamine B (SRB) assay at 2, 5, 9 and 14 days after UVC illumination (**Figure 1A**). 254-nm and 277-nm illumination resulted in low cell viability throughout the duration, while 222-nm far UVC caused the cell viability to be significantly lower than the non-illuminated control cells at days 9 and 14. The UVC-induced lowering of cell viability is dose-dependent, as evidenced by the SRB assay taken at 9 days after UVC illumination (**Figure 1B**).

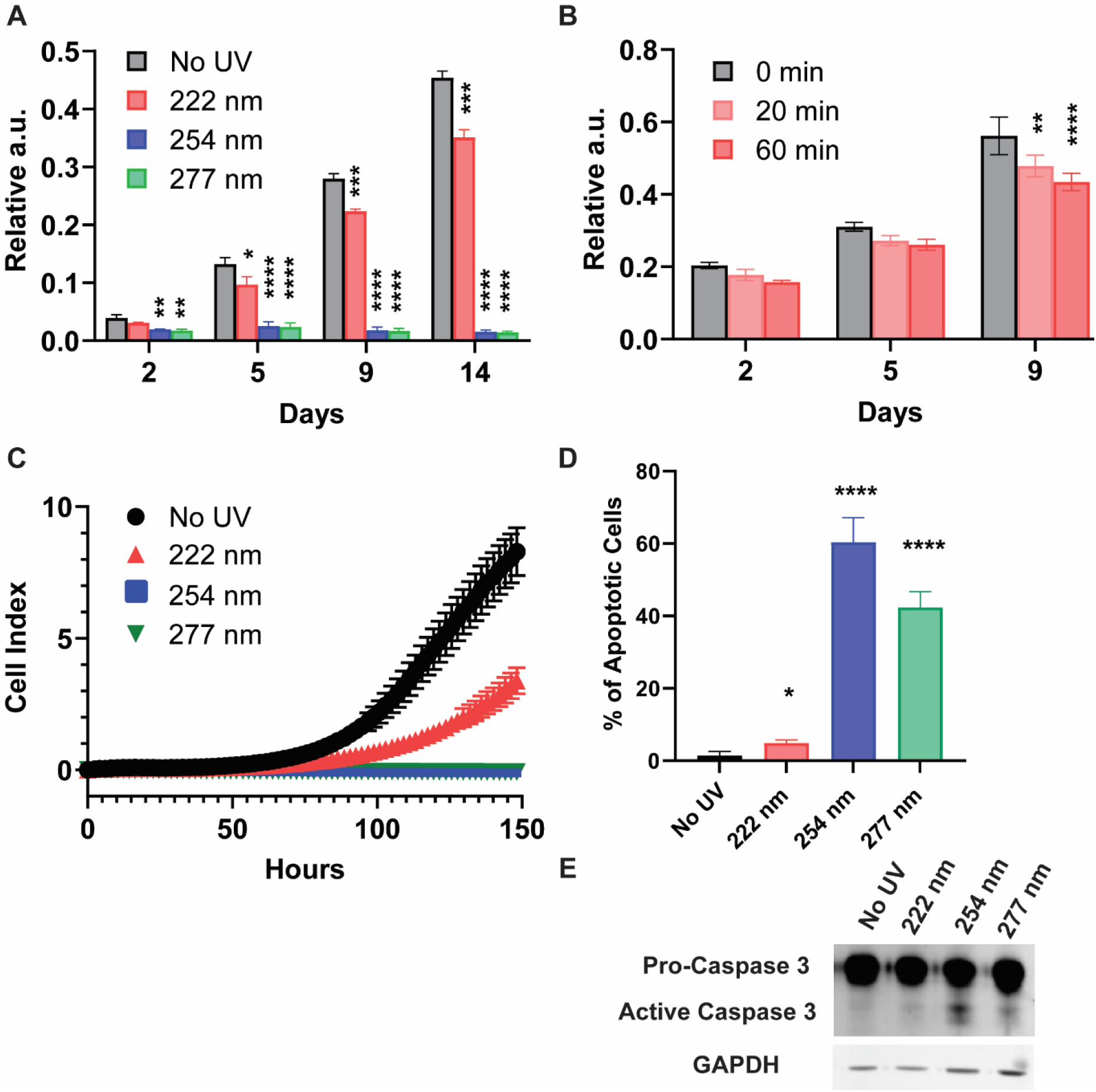
(a) SRB assay for ARPE-19 cells exposed to either no UVC or 60 minutes of respective UVC wavelengths and then incubated for the indicated number of days. Values are reported as mean +/- S.D. from n = 3 experiments. (b) SRB assay for ARPE-19 cells exposed to either 0, 20 or 60 minutes of 222-nm far UVC and then incubated for the indicated number of days. Values are reported as mean +/- S.D. from n = 3 experiments. (c) Dynamic monitoring of cell numbers through xCelligence platform. Values are reported as mean +/- SD from n = 8 replicates. (d) Trypan blue staining assay performed on ARPE-19 cells exposed to either no UVC or 60 minutes of respective UVC wavelengths and then incubated for 1 day. Values are reported as mean +/- SD from n = 4 experiments. (e) Representative Western blot analysis of pro-caspase 3 versus cleaved caspase 3 after either no UVC or 60 minutes of respective UVC wavelengths. GAPDH immunoblotting is used as an internal control.

To gain a comprehensive and dynamic understanding of the cellular growth profile, we utilized the xCELLigence platform to track the number of cells per condition. In agreement with the results from SRB assays, 222-nm UVC results in significant reduction of cell numbers while 254-nm and 277-nm UVC saw no viable cell growth within the 6-day experimental period (**Figure 1C**). The 222nm-induced reduction of cell viability is dose-dependent as demonstrated in **Figure S1**.

We then carry out trypan blue staining to track the cell death of ARPE19 1 day after varied UVC wavelength illumination (**Figure 1D**), where 254-nm and 277-nm UVC posted a high percentage of trypan blue stained cells at 60.5% and 41.2% respectively. This is corroborated by Western blot analysis which saw caspase-3 activation prominently in 254-nm- and 277-nm-lit cells. (**Figure 1E**)

### Differential signalling pathway activation under varied UVC wavelengths

The mitogen-activated protein kinases (MAPKs) have been well established to mediate the UV response, which determine the cell fate^9^. These include the extracellular signal-regulated kinases (ERKs), the c-Jun NH2-terminal kinases (JNKs) and the p38 kinases (**Figure 2A**). We hypothesize that the MAPKs could be regulated differently by the UVC wavelengths, which result in such diverse cellular outcomes. As such, we subjected ARPE-19 cells to 20 and 60 minutes of the respective UVC wavelengths and monitored the activation of the MAPKs. **Figure 2B** shows that while the ERK and p38 pathways are activated under all three UVC wavelengths, phosphorylation of JNK/ATF2 pathway is not observed under 222-nm illumination.

**Figure 2:**
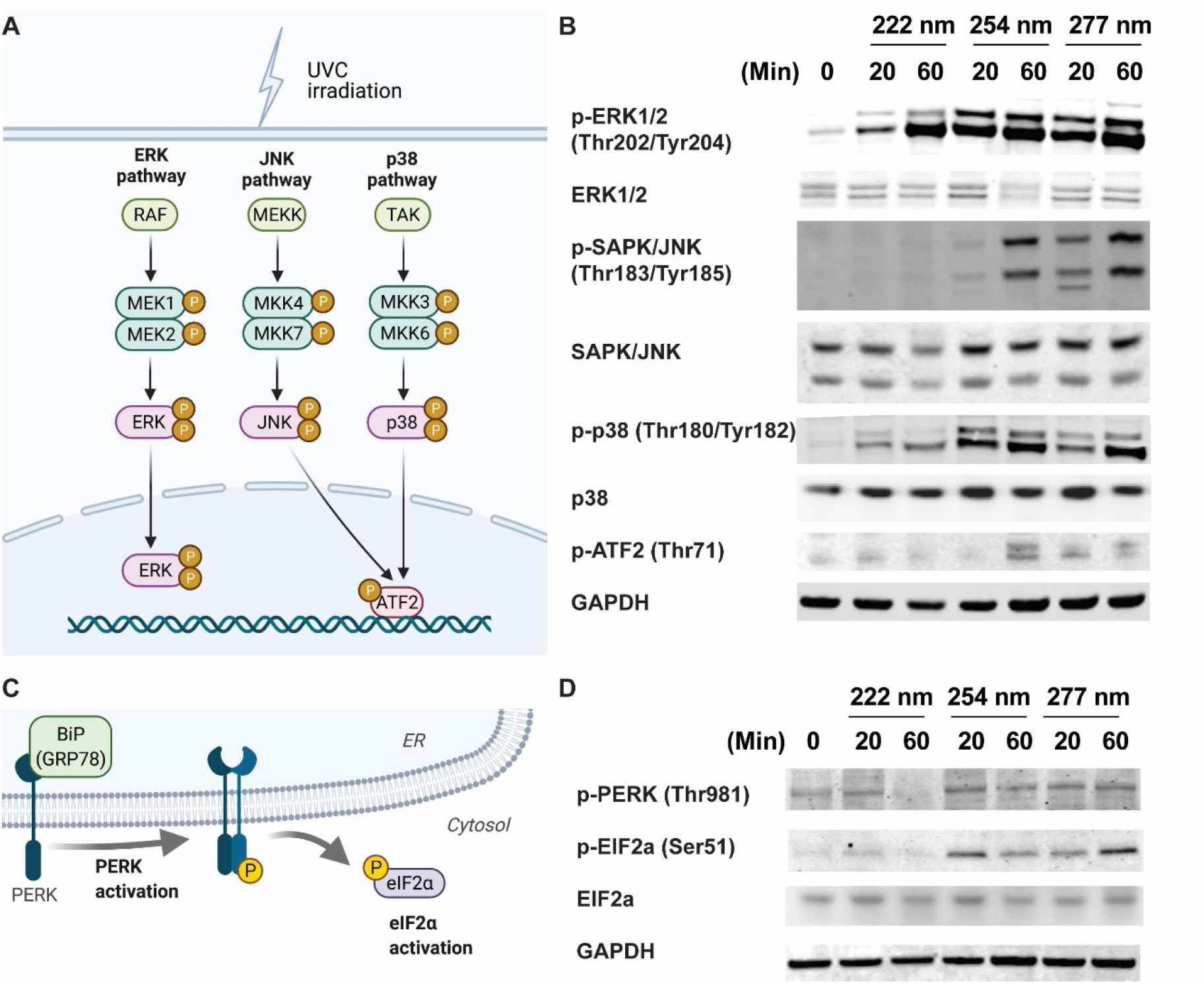
Differential activation of MAPK and ER-stress signalling pathways results in the differential activation of apoptotic pathways. (a) Schematic indicating the classical MAPK signalling pathways studied due to classical 254-nm UVC irradiation. (b) Representative Western blots of phosphor-ERK (Thr202, Tyr204), total ERK1/2, phosphor-JNK/SAPK (Thr183, Tyr185), total JNK1/2, phosphor-p38 (Thr180, Tyr182), p38, phosphor-ATF2 (Thr69) and GAPDH from ARPE-19 cells exposed to 0, 20 and 60 minutes of respective UVC wavelengths. (c) Schematic indicating the proteins involved in endoplasmic reticulum-induced stress pathway. (d) Representative Western blots of phosphor-PERK (Thr980), total ERK1/2, phosphor-EIF2a (Ser49), EIF2a and GAPDH from ARPE-19 cells exposed to 0, 20 and 60 minutes of respective UVC wavelengths.

It has also been known that the PERK/eIF2a axis is crucial for mediating endoplasmic reticulum-stress signalling upon UV irradiation^18^ (**Figure 2C**). We noticed that the phosphorylation events of PERK and eIF2a did not occur in 222-nm-lit cells as compared to 254-nm and 277-nm-illuminated cells (**Figure 2D**). Given that both JNK and PERK/eIF2a pathway activation have been linked to apoptosis events, it could be inferred that the lack of activation of these stress signalling pathways in 222-nm-lit cells could be responsible for sustaining the ARPE19 cell survival^19,20^.

### DNA-associated damage or associated repair mechanisms is observed in 222-nm-lit cells

Upon understanding the lack of apoptotic mechanisms in 222-nm-lit cells, we next asked if physical damages are sustained on these cells as compared to other wavelengths. Using Coomassie staining, we saw that ARPE19 cells subject to 60 minutes of 222-nm treatment had reduction in intensity of several protein bands (**Figure S2A**). 4-hydroxyonenal (4-HNE) staining further revealed increases in oxidative stress due to increased lipid peroxidation chain reaction (**Figure S2B**).

UVC has also been well known to cause DNA damage to cells, where the formation of cyclobutane dimers and (6,4)-photoproducts have been well characterized^2,8^. In recent years, yH2AX has emerged as a reliable marker for double strand breaks and their repair^21^. We hypothesize that DNA in mammalian cells could be potentially underdoing damages under the different UVC wavelengths and the subsequent DNA-associated damages and repair mechanisms could be visualized with yH2AX staining. Using fluorescence microscopy, we observed that 222-nm far UVC resulted in the highest percentage of yH2AX positive cells compared to 254-nm and 277-nm UVC wavelength illumination. (**Figure 3A-B**) The number of foci in 222-nm illuminated cells remains high 48 hours after the initial illumination, indicating the persistence of such foci. (**Figure 3C**)

**Figure 3:**
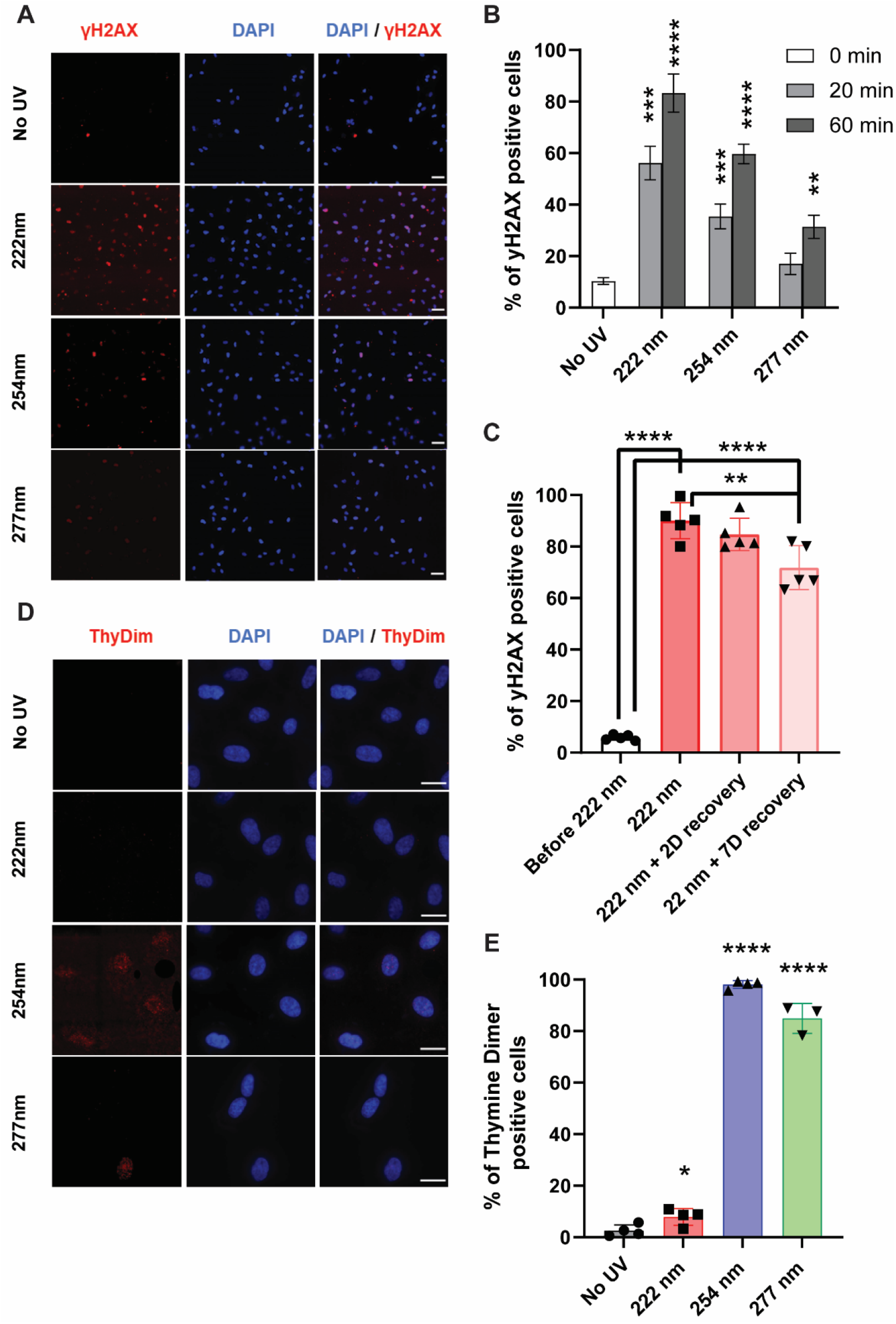
222-nm far UVC results in significant DNA damage which does not include typical UVC-induced DNA cyclobutane dimer formation. (a) γH2AX staining in ARPE19 cells upon 60 minutes of respective UVC irradiation. Scale bars = 50 μm. (b) Quantification of the extent of γH2AX activation in the ARPE19 cells upon 0, 20 and 60 minutes of UVC illumination. Values are reported as mean +/- SD from n = 3 experiments. (c) Quantification of the extent of γH2AX activation in the ARPE19 cells exposed to 60 minutes of 222-nm far UVC illumination and then recovered 0, 2 and 7 days after the far UVC treatment. Values are reported as mean +/- SD from n = 5 experiments. (d) Thymine dimer staining in ARPE19 cells upon 60 minutes of respective UVC irradiation. Scale bars = 20 μm. (e) Quantification of the extent of thymine dimer formation in the ARPE19 cells upon 60 minutes of UVC illumination. Values are reported as mean +/- SD from n = 3-4 experiments.

In comparison, 222-nm far UVC did not elicit any cyclobutane dimer formation while the formation of these dimers is most apparent due to 254-nm and 277-nm illumination. (**Figure 3D-E**) This indicates that the DNA damages or repair mechanisms elicited from 222-nm far UVC do not come from cyclobutane dimer formation and potentially derived from other forms of stress insults. These observations are congruent in HEKA keratinocytes (**Figure S3**), which indicates similar mechanisms could be at play in different mammalian cells.

### Perturbation of transcription and cellular signaling events is observed in 222-nm-lit cells

Given that 222-nm far UVC illumination poses damages on the DNA of ARPE-19 cells, we next investigated whether the cellular transcriptional programs and key signaling modalities are restructured. We first examined transcriptional changes via qRT-PCR of ARPE-19 cells 1 day after 60 minutes of 222-nm far UVC illumination and focused on tumor suppressor genes (Rb1 and P53), cancer-related genes (SNAI1, E2F1, E2F2, CDK4) and stress-related genes (BMP4, NRF2, p21, IL6 and HIF1A). (**Figure 4A**) Notably, there is a significant upregulation of SNAI1 and p21 genes, and key genes that are downregulated include BMP4, p53, E2F2 and E2F1 genes. These are corroborated by Western blot analyses (**Figure 4B-C**) where both tumor suppressor genes Rb1 and p53 saw a reduction in band intensity in 1-3 days after 222-nm far UVC illumination, whilst phosphor-ERK and its downstream signaling target SNAI1 posted increase 1-3 days after 222-nm exposure. Interestingly, the protein levels of the tumor suppressor genes and the extent of phosphor-ERK recovered to the original levels at 6 days after the exposure.

**Figure 4:**
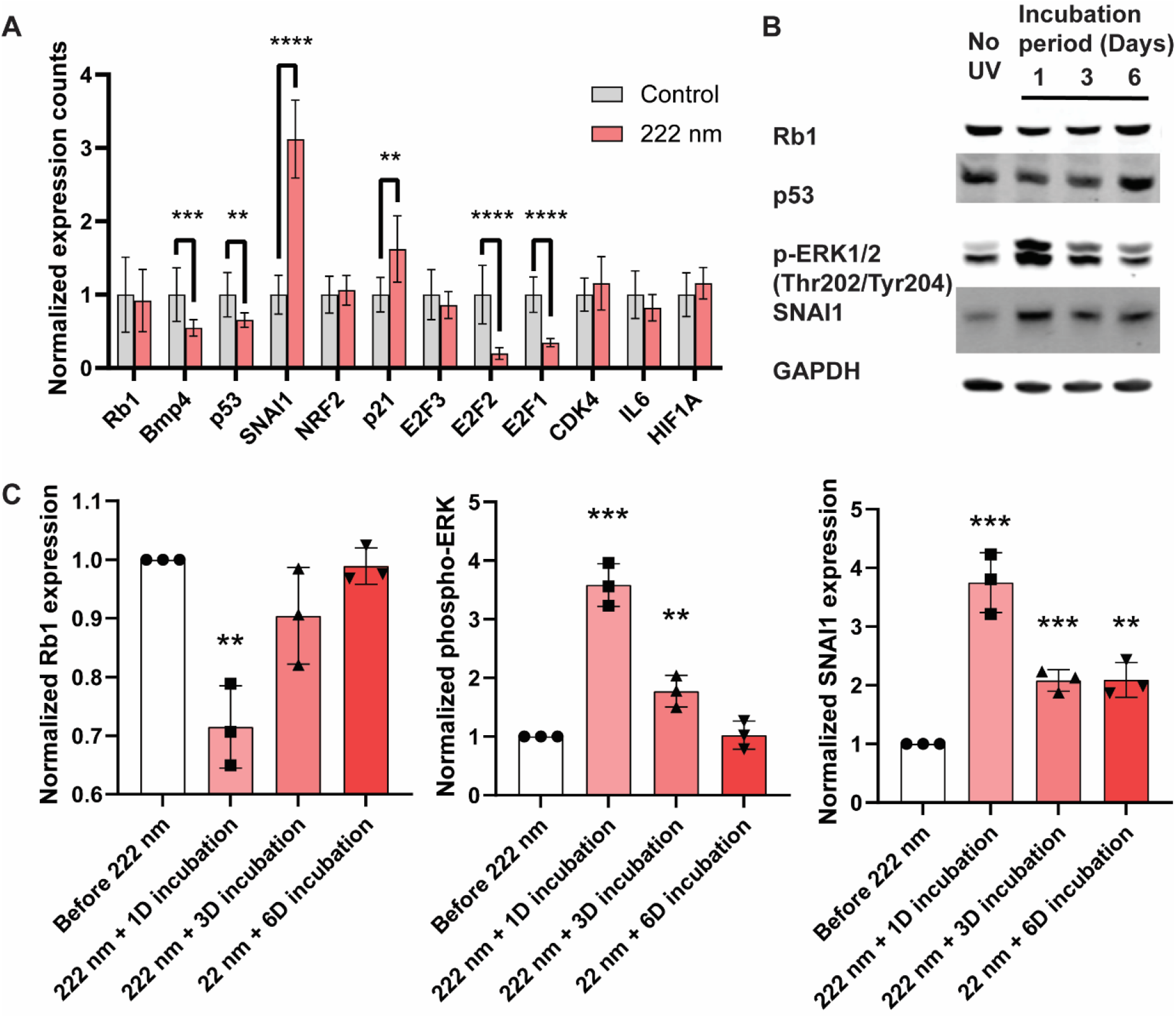
RT-PCR and Western blot analysis of ARPE-19 cells after 222-nm far UVC illumination. (a) RT-PCR analysis of selected genes 1 day after 60 minutes of 222-nm far UVC illumination. (b) Representative Western blots of Rb1, p53, phosphor-ERK (Thr202, Tyr204), SNAI1 and GAPDH from ARPE-19 cells 1, 3 and 6 days after being exposed to 60 minutes of 222-nm far UVC illumination. (c) Quantification of the band intensity for Rb1, p53, phosphor-ERK (Thr202, Tyr204) and SNAI1 relative to internal control GAPDH in the ARPE19 cells 1, 3 and 6 days upon 60 minutes of UVC illumination. Values are reported as mean +/- SD from n = 3 experiments.

### Analysis of global RNA transcripts one week after 222-nm exposure

In order to investigate whether the changes in transcriptional activity is a transient phenomenon, we collected RNA from cells one week exposed to no light or 222-nm UVC far illumination and performed RNA sequencing on both sets. **Figure 5A** plots the top differentially expressed genes as derived from the RNA sequencing results and **Figure S4** depicts the principal component analysis of the triplicates used in the analysis. We then picked out the statistically significant genes (p < 0.05, n = 2361) and analyzed this subset using Ingenuity Pathway Analysis (IPA). **Figure 5B** reveals the top ingenuity canonical pathways where top upregulated pathways include DNA damage, checkpoint control and regulation while top downregulated pathways include cell cycle and replication pathways. The same trends are reflected in the Diseases and Disorders set where a large percentage (∼90%) of the genes are implicated in the larger category of Cancer and Organismal Injury and Abnormalities. (**Figure 5C**) Out of these pathways, the top cellular functions cover on cell death and survival and cell cycle. We then summarized these IPA analysis findings into the summary figure (**Figure 5D**) which shows the top pathways implicated include formation of gamma H2AX foci and senescence of cells being upregulated whilst cell growth related pathways such as entry into interphase and colony formation being affected. **Files S1, S2 and S3** contain further information regarding the top differentially expressed genes, top regulator pathway and analyzed upstream regulator networks as informed by the IPA analysis.

**Figure 5:**
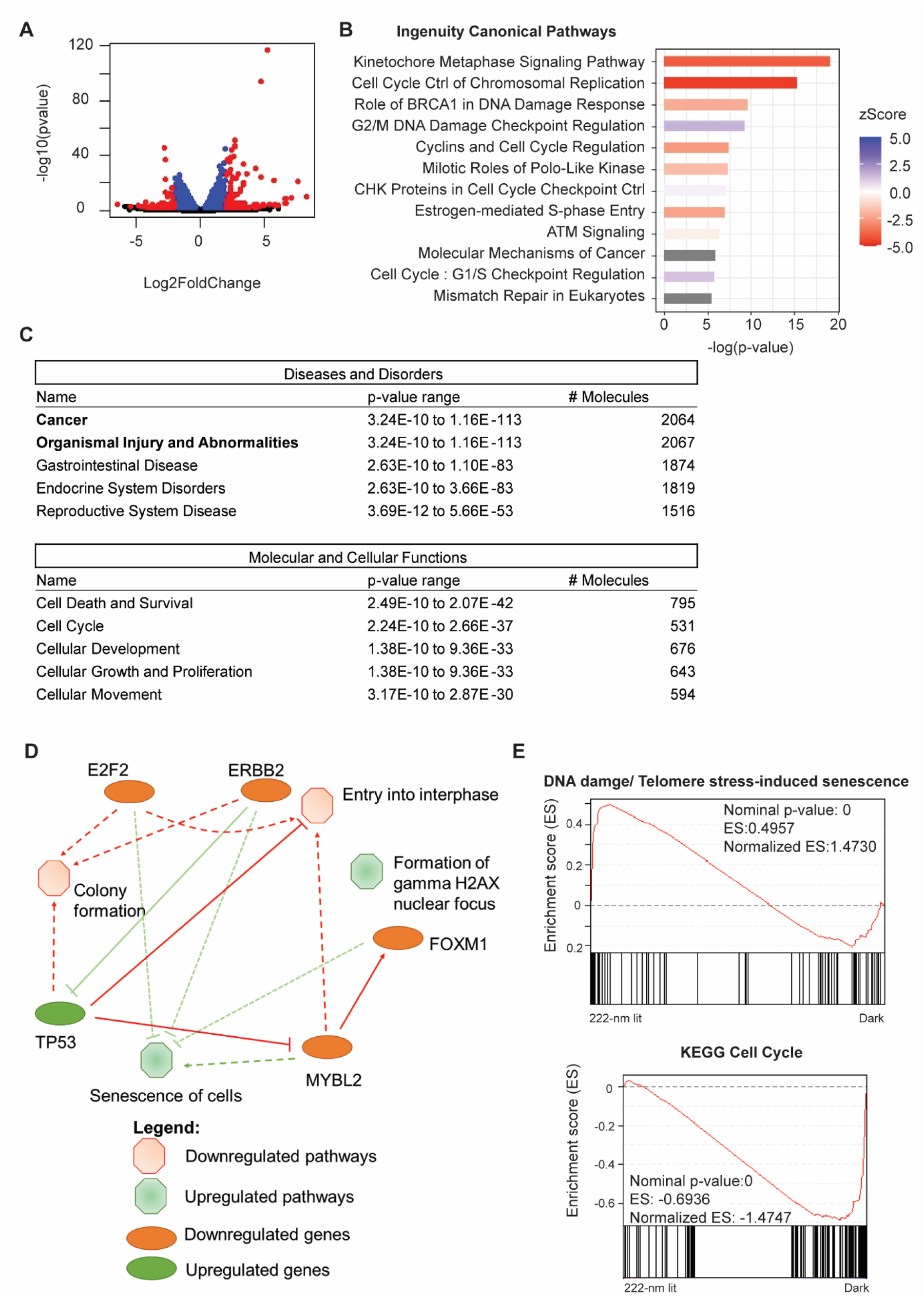
RNA sequencing analysis of global RNA transcriptomic changes one week after 222-nm far UVC illumination. (a) Volcano plot of statistically significant differentially expressed genes at p < 0.05. Highlighted in red are genes that are either upregulated with log2FoldChange at more than 2 and p < 0.05 (n = 140), or downregulated with log2FoldChange at less than 2 and p < 0.05 (n = 73). (b) Top ingenuity canonical pathways involved due to 222-nm illumination with ingenuity pathway analysis performed on 2361 statistically significant differentially expressed genes at p < 0.05. A positive activation zScore reflects upregulation of pathway in 222-nm illuminated ARPE-19 cells while a negative activation zScore reflects the opposite. (c) Top diseases and disorders and molecular and cellular function categories derived from ingenuity pathway analysis out of 2361 statistically significant differentially expressed genes. (d) Ingenuity pathway analysis graphical summary providing a quick overview of the major biological themes and how they relate to each other. (e) Gene Set Enrichment analysis of all 20358 genes detected in the RNA sequencing found upregulation of DNA damage and telomere-stress induced senescence pathway and a reduction of cell cycle genes in 222-nm illuminated ARPE-19 cells.

In addition, we also conducted Gene Set Enrichment Analysis (GSEA) based on 20358 genes being detected in the RNA sequencing profiles, and found that the results corroborated with the IPA analysis. **Figure 5E** shows the two representative key pathways up/downregulated in the form of DNA damage/Telomere stress-induced senescence (Normalized Enrichment Score of 1.473) and KEGG Cell Cycle (Normalized Enrichment Score of −1.4747). Further GSEA analysis could be found in **Figure S5**.

## Discussion

As the COVID-19 pandemic strikes and highlights the importance of public disinfection tools in coping with COVID-19 and future viruses in the post-COVID world, 222-nm far UVC has emerged as a potential safe solution in disinfecting viruses, including SARS-CoV-2, whilst presenting as a safe option that allows for human exposure^12–14,22,23^. The safety presumption of 222-nm far UVC has been largely based on the argument that the 222-nm far UVC cannot penetrate the outer, non-living cells of the eyes and skin, and that 222-nm far UVC does not induce typical UVC-induced DNA lesions in human keratinocyte model and skin of exposed hairless mice. The former has yet been validated experimentally and to date, there has been a lack of long-term studies on human exposure. While data presented by the latter seems promising, the results call for deeper and further investigations as they only covered the typical UVC-induced DNA lesions and not general classes of DNA damages.

In this paper, we exposed the skin and eye cells to 20 and 60 minutes of respective UVC wavelength and studied the immediate and longer term effects of UVC damage towards cellular health and viability. In agreement with earlier reports, 222-nm lit cells survived and retained ability to grow without activation of apoptosis whilst 254-nm and 277-nm lit cells showed decreased viability and had apoptosis activated^24^. We then studied the mechanisms leading to the lack of such activation and found that potentially pJNK/ATF2 and the endoplasmic reticulum stress pathways may be involved. However, we found that the 222-nm lit cells had a lower growth rate compared to unilluminated controls. We then pursued the DNA damage pathways and saw that while 222-nm far UVC did not incur any thymine dimer formation unlike 254-nm UVC lamp and 277-nm UVC LED, it recorded the highest incidence of γH2AX nuclear foci indicating DNA damage. This was particularly alarming given that the tumor suppressor genes, Rb1 and p53, are downregulated in the 222-nm lit cells one to three days after the exposure. RNA sequencing of cells exposed to 222-nm illumination was also conducted to find that the cellular survival and cell cycle pathways are re-sculpted with the summary analysis showing the upregulation of γH2AX nuclear foci and senescence of cells as corroborating with all the previous results.

It is alarming that clinical scientists and researchers have taken the initial optimism regarding the safety profile of 222-nm far UVC and started to pursue clinical trials for performing such illumination on open wounds^25^ and in healthy humans^26^. The excitement generated in the field has also seen huge interest for this technology in hospitals and facilities management adopting the tool for open human exposure.

## Limitations of Study

In this study, we did not extend the impact of different UVC wavelengths on cells grown in a three-dimensional model or organism level. Whilst the study is limited in only in vitro studies, we would express our hope for further in-depth studies on the molecular changes that 222-nm far UVC would cause and be more careful in widespread adoption.

## Associated Content

**Supporting Information**. The following files are available free of charge.

**Figure S1:** Dose-dependence curves of ARPE-19 cells subjected to no UV, 20 minutes of 222-nm illumination and 1 hour of 222-nm illumination.

**Figure S2:** Different UVC wavelength result in varied levels of protein and lipid damage.

**Figure S3:** γH2AX and thymine dimer staining in HEK-A cells upon 60 minutes of respective UVC irradiation. Scale bars = 50 μm.

**Figure S4:** Principal component analysis of RNA sequencing results.

**Figure S5:** Further GSEA analysis including upregulated and downregulated pathways in 222-nm lit cells.

**File S1:** Top differentially expressed genes between 222-nm lit versus unlit cells.

**File S2:** Top regulator pathways as predicted by IPA analysis.

**File S3:** Top predicted changes in upstream regulators from IPA analysis.

## Acknowledgments

We thank Dr. Shawn Tan and members of the HWP lab for useful suggestions and comments. We also thank Dr. Yuanjie Liu for providing technical advice and help.

## Funding

National Research Foundation grant NRF2020NRF-CG002-035 (QO, JWRT)

Accelerate Technologies Gap-funded project GAP/2020/00392 (JWRT, QO)

## Author contributions

QO, JWRT, WH conceptualized the study. QO and JWRT designed the experiments. QO, WW and JDC performed the experiments. QO and WH wrote the manuscript.

## Competing interests

Authors declare that they have no competing interests.

## Materials and Methods

### KEY RESOURCES TABLE

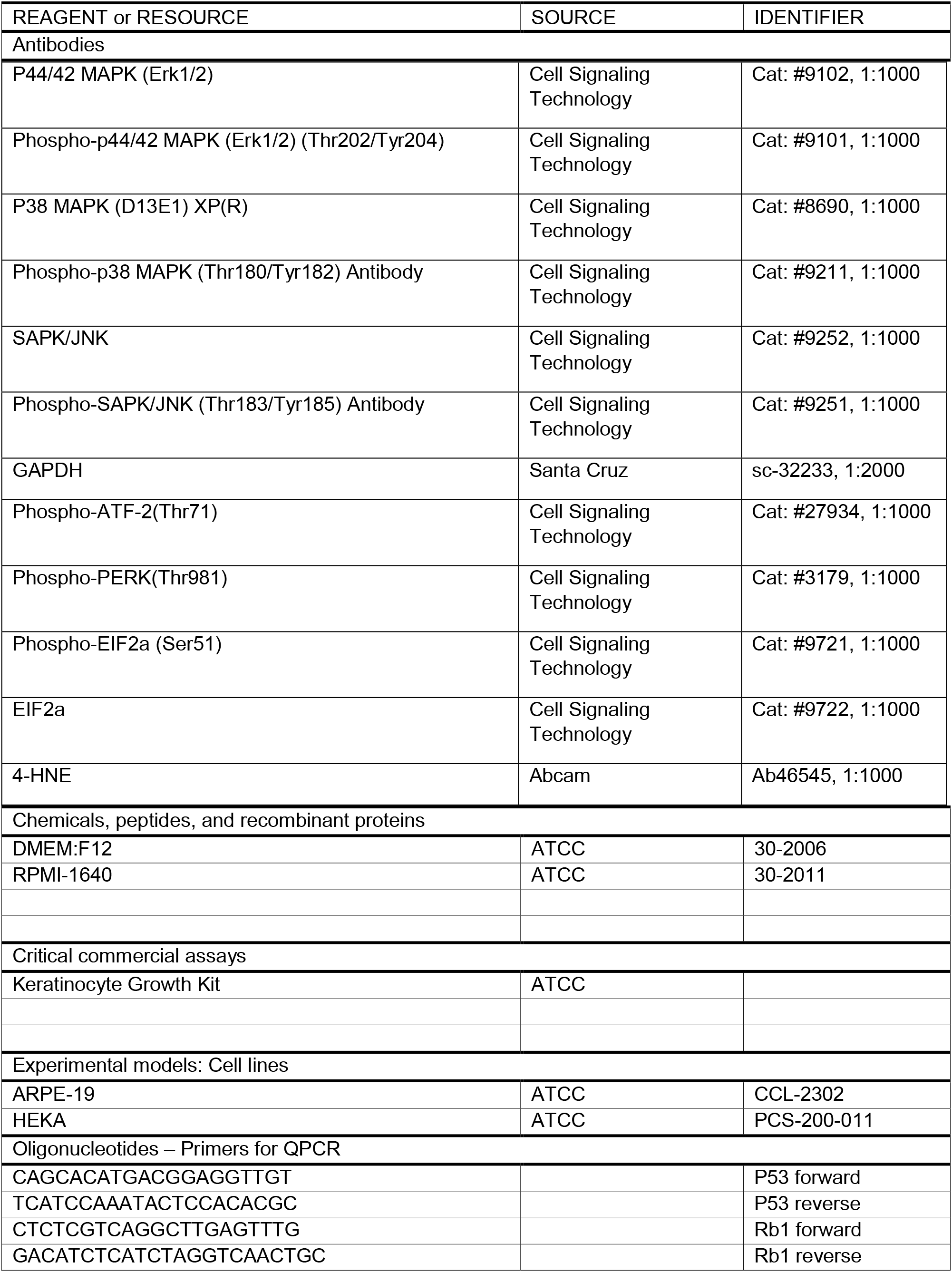
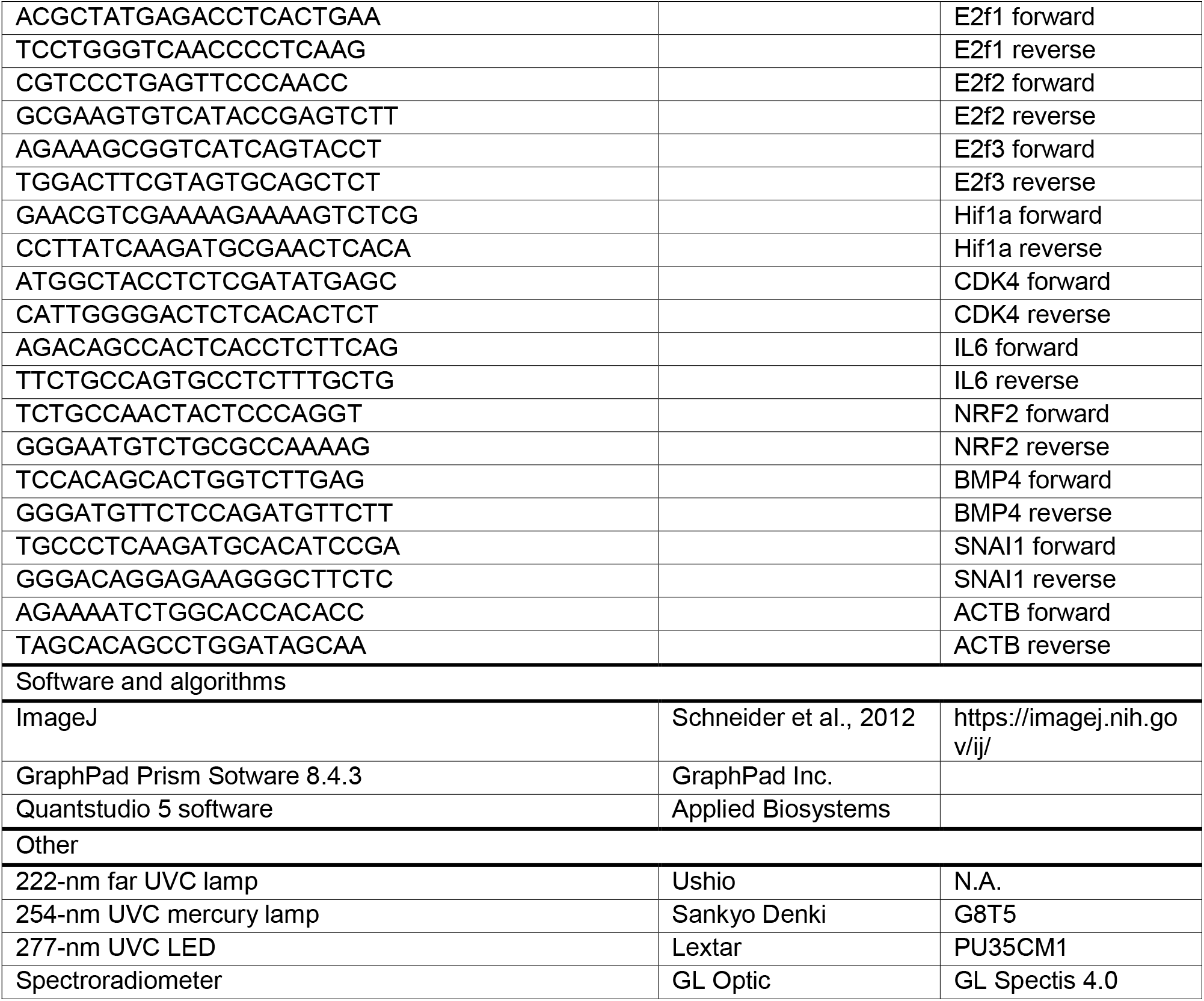

#### UVC sources and irradiance measurements

To understand the effect of UVC wavelength on human cell lines, three different UVC light sources – 222 nm far UVC lamp (Ushio), 254 nm UVC mercury lamp (Osram) and 277 nm UVC LED (Lextar PU35CM1) were used in this study. The detailed method has been described in an earlier work.^15^ These UVC light sources were measured using a calibrated spectroradiometer (GL Spectis 4.0) with an absolute measurement uncertainty of less than 6% from 200 to 500 nm. To provide a comparative UVGI efficacy study between these UVC light sources, the radiant intensity of the far UVC and mercury lamp is measured at different distances while the UVC LED is driven at different constant drive currents to obtain a common UV intensity of 73 μW/cm^2^. The UVC LED, with a beam angle of 120°, is assembled into a 5 × 5 array at a working distance of 12 cm to ensure uniform UV intensity across the surface of the petri dish.

#### Cell culture

The ARPE-19 retinal cells were grown in DMEM:F12 (ATCC Catalog No. 30-2006) supplemented with 10% Fetal Bovine Serum (FBS), 100 U/ml penicillin and 100 μg/ml streptomycin (Sigma-Aldrich, St. Louis, MO). The HEK-A keratinocytes (ATCC PCS-200-011) were cultured in Dermal Cell Basal Medium supplemented with the keratinocyte growth kit (ATCC), 100 U/ml penicillin and 100 ug/ml streptomycin (Sigma-Aldrich, St. Louis, MO).

#### Cell viability assays

To measure the effect of different UVC illumination on cell viability, the cells were exposed to 0.073 μW/cm^2^ of respective wavelength for an hour and then analyzed after the defined recovery time. These are performed with the xCELLigence platform (Roche) and also through sulforhodamine B (SRB) assay.

##### xCELLigence platform

The cells were irradiated in 100mm petri dishes, allowed to recover overnight before being seeded into the appropriate 96-well xCELLigence plates at a density of 1000 cells per well. Following seeding, the cells were monitored every 10 minutes by the xCELLigence system (Roche) for proliferation, attachment, and spreading. The impedance detection was performed for a total of 6 days. Real-Time Cell Analysis 2.0 (Roche) software was used to analyze the data.

##### SRB assay

The viability of each condition was determined using a colorimetric SRB assay. ^16^ The cells were irradiated in 100mm petri dishes, allowed to recover overnight before being seeded into 96-well tissue culture plates at a density of 1000 cells per well. For the defined number of days following irradiation, 100 μl cold 10% trichloroacetic acid added and allowed to incubate at 4°C for 1 hour. The plates are then washed with distilled water, dried in 37°C for 30 minutes before 80 μl of SRB dye is added to each well for 5 minutes. The plates are then washed with 1% acetic acid, dried in 37°C for 30 minutes and the dye was then resolubilized in 10 mM Tris at room temperature for an hour. The plates are then recorded for the absorbance at a wavelength of 510 nm, with the values expressed in arbitrary units. All SRB assay experiments were performed in triplicate.

#### Immunofluorescence experiments

To assess whether various UVC illumination causes different types of DNA damages on ARPE-19 and HEK-A cells, immunostaining was performed to detect the presence of γH2AX and thymine dimers in these cells. Briefly, 2 × 10^5^ cells were plated in each petri dish one day before the experiment. After the described duration of UVC illumination, the cells were washed with PBS and fixed with 4% paraformaldehyde at room temperature for 15 minutes and washed with PBS before being labelled with anti-phospho-Histone H2A.X (Ser139) antibody (Sigma Aldrich, Cat#:05-636) 1:500 or anti-Thymine dimer antibody (Novus Biological, Cat#: NB600-1141) 1:250 in PBS containing 2% bovine serum albumin (BSA) and 0.1% TBS-T. Cells were then washed with PBS and labelled with goat anti-rabbit Alexa Fluor-568 (Life Technologies, Grand Island, NY) in PBS containing 2% BSA at room temperature for an hour with gentle shaking. Following washing with PBS, the cells were stained with DAPI and observed with the 40x objective of Nikon Ti-2 TIRF microscope.

#### Western blot experiments

All samples for Western blots were lysed in NP40 buffer supplemented with phosphatase and protease inhibitors. Approximately 10 μg of protein sample is loaded for western blot experiments. The lysates were then subjected to SDS gel electrophoresis before being transferred to nitrocellulose membranes using iBlot2 (Life Technologies), blocked with 5% BSA in TBS with 0.1% Tween-20 and incubated with primary antibodies. Membranes were then incubated with rabbit-IRDye 800 CW secondary antibodies and imaged on an Odyssey CLx (LI-COR).

#### RNA extraction and analysis

RNA was isolated from ARPE-19 cells by Trizol isolation and genomic DNA was subsequently removed with DNAseI (Invitrogen). cDNA was then generated from 1 μg of RNA using RevertAid First Strand cDNA Synthesis Kit (Thermofisher) as per manufacturer’s recommendations. Quantitative PCR was then carried out with PowerUp SYBR Green Master Mix (Life Technologies) and the analysis was performed using the Quantstudio 5 software.

#### RNA sequencing

Total RNA from control and 222nm-treated ARPE-19 cells was isolated and 1 μg of RNA per sample was used as the input material. Next generation sequencing library preparations were constructed according to the manufacturer’s protocol. The poly(A) mRNA isolation was performed using Poly(A) mRNA Magnetic Isolation Module or rRNA removal Kit. The mRNA fragmentation and priming were performed using First Strand Synthesis Reaction Buffer and Random Primers. First strand cDNA was synthesized using ProtoScript II Reverse Transcriptase and the second-strand cDNA was synthesized using Second Strand Synthesis Enzyme Mix. The purified double-stranded cDNA by beads was then treated with End Prep Enzyme Mix to repair both ends and add a dA-tailing in one reaction, followed by a T-A ligation to add adaptors to both ends. Size selection of Adaptor-ligated DNA was then performed using beads, and fragments of ∼400 bp (with the approximate insert size of 300 bp) were recovered. Each sample was then amplified by PCR using P5 and P7 primers, with both primers carrying sequences which can anneal with flow cell to perform bridge PCR and P5/ P7 primer carrying index allowing for multiplexing. The PCR products were cleaned up using beads, validated using an Qsep100 (Bioptic, Taiwan, China), and quantified by Qubit3.0 Fluorometer (Invitrogen, Carlsbad, CA, USA). Then libraries with different indices were multiplexed and loaded on an Illumina Novaseq instrument according to manufacturer’s instructions (Illumina, San Diego, CA, USA). Sequencing was carried out using a 2×150 paired-end (PE) configuration.

#### Quantification of gene expression

Quality of the sequences was assessed with FastQC v0.11.5^17^. No further data filtering and trimming were performed. The paired FASTQ files were aligned to hg19 reference genome using GENCODE v.36 gene annotations and STAR v2.6.0a splice aware aligner. Unique transcripts were then assembled from merged alignment files, and abundance of the transcripts are generated by featureCounts. Differentially expressed genes (DEGs) were determined with DESeq2.

#### Pathway analyses

In total, 2359 genes were differentially expressed between the illuminated and control group (adjusted Benjamini-Hochberg p-value < 0.05), with a total of 1319 down-regulated and 1040 upregulated genes. These genes were used for Ingenuity Pathway Analysis (Qiagen). For Gene Set Enrichment Analysis, it was performed with the software version 4.0.3.

## Notes

### Competing Interest Statement

The authors have declared no competing interest.

