## Supplementary Info for "222-nm far UVC exposure results in DNA damage and transcriptional changes to mammalian cells"

**Supplemental Figure Legends**

**Figure S1:** Dose-dependence curves of ARPE-19 cells subjected to no UV, 20 minutes of 222-nm illumination and 1 hour of 222-nm illumination. (c)  Dynamic monitoring of cell numbers through xCelligence platform. Values are reported as mean +/- SD from n = 8 replicates.

**Figure S2:** Different UVC wavelength result in varied levels of protein and lipid damage. **(A)** Coomassie staining of protein lysates of ARPE-19 cells subject to 0, 10, 30 and 60 minutes of respective UVC wavelength. **(B)** Western blot analysis of 4-HNE adducts was performed on protein lysates of ARPE-19 cells subject to 0, 20 and 60 minutes of respective UVC wavelength.

**Figure S3:** **(A)** γH2AX staining in HEK-A cells upon 60 minutes of respective UVC irradiation. Scale bars = 50 µm. Quantification of the extent of γH2AX activation in the HEK-A cells upon 0, 20 and 60 minutes of UVC illumination. Values are reported as mean +/- SD from n = 3 experiments. **(B)** Thymine dimer staining in HEK-A cells upon 60 minutes of respective UVC irradiation. Scale bars = 20 µm. Quantification of the extent of thymine dimer formation in the HEK-A cells upon 60 minutes of UVC illumination. Values are reported as mean +/- SD from n = 3 experiments.

**Figure S4:** Principal component analysis of RNA sequencing results.

**Figure S5:** Further GSEA analysis including upregulated pathways in 222-nm lit cells in **(A)** Epithelial-mesenchymal transition, **(B)** Angiogenesis, **(C)** Integrin-cell surface interaction and downregulated pathways such as **(D)** E2F targets.

**Supplemental Files**

**File S1:** Top differentially expressed genes between 222-nm lit versus unlit cells.

**File S2:** Top regulator pathways as predicted by IPA analysis.

**File S3:** Top predicted changes in upstream regulators from IPA analysis.

**Supplementary Figures**

**Figure S1**

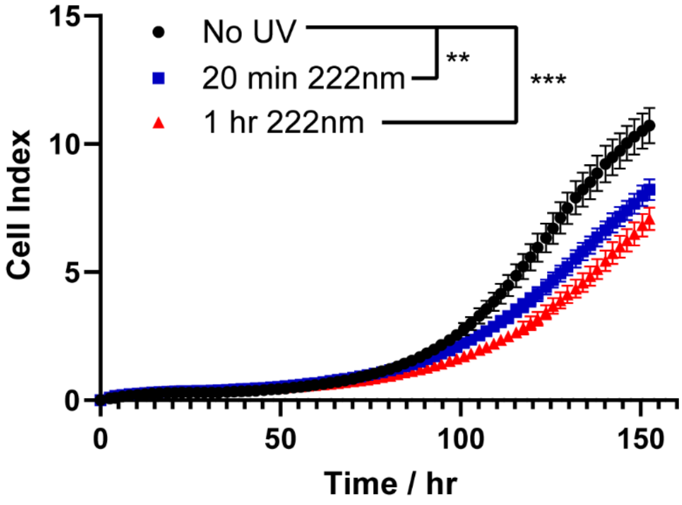

**Figure S2**

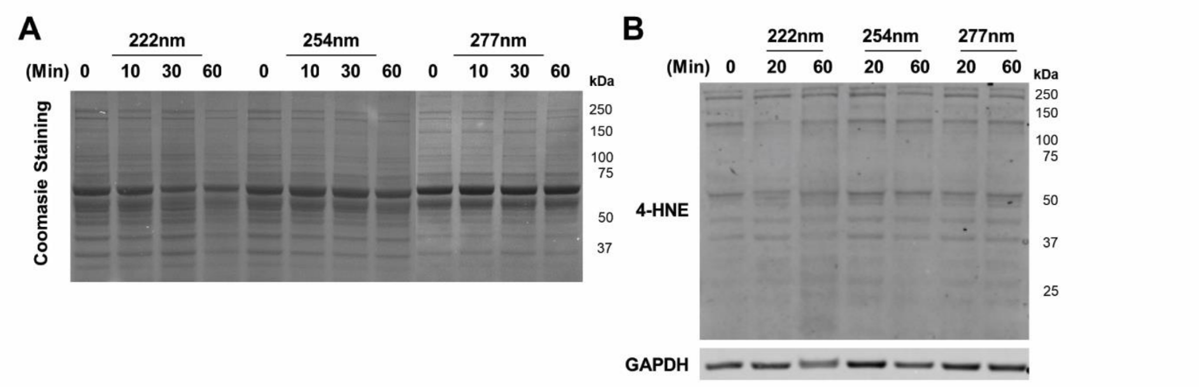
A
222nm
254nm
277nm
(Min) 0 10 30 60 0 10 30 60 0 10 30 60
kDa
250
150
100
75
50
37
Coomasie Staining
B
222nm
254nm
277nm
(Min) 0 20 60 20 60 20 60
kDa
250
150
100
75
50
37
25
4-HNE
GAPDH

**Figure S3**

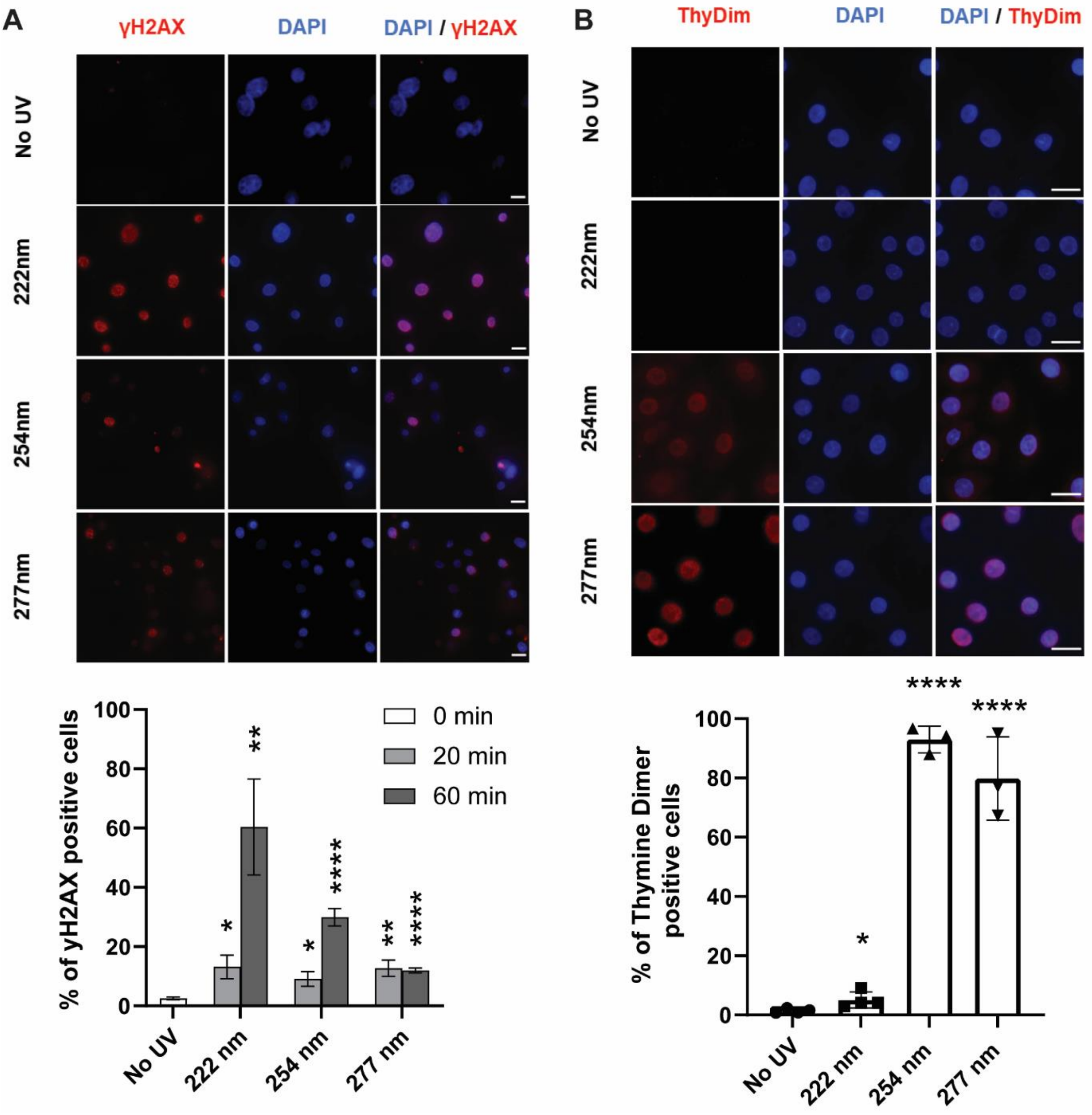

A
γH2AX
DAPI
DAPI / γH2AX
No UV
222nm
254nm
277nm
% of γH2AX positive cells
0 min
20 min
60 min
No UV
222 nm
254 nm
277 nm
B
ThyDim
DAPI
DAPI / ThyDim
No UV
222nm
254nm
277nm
% of Thymine Dimer positive cells
No UV
222 nm
254 nm
277 nm

**Figure S4**

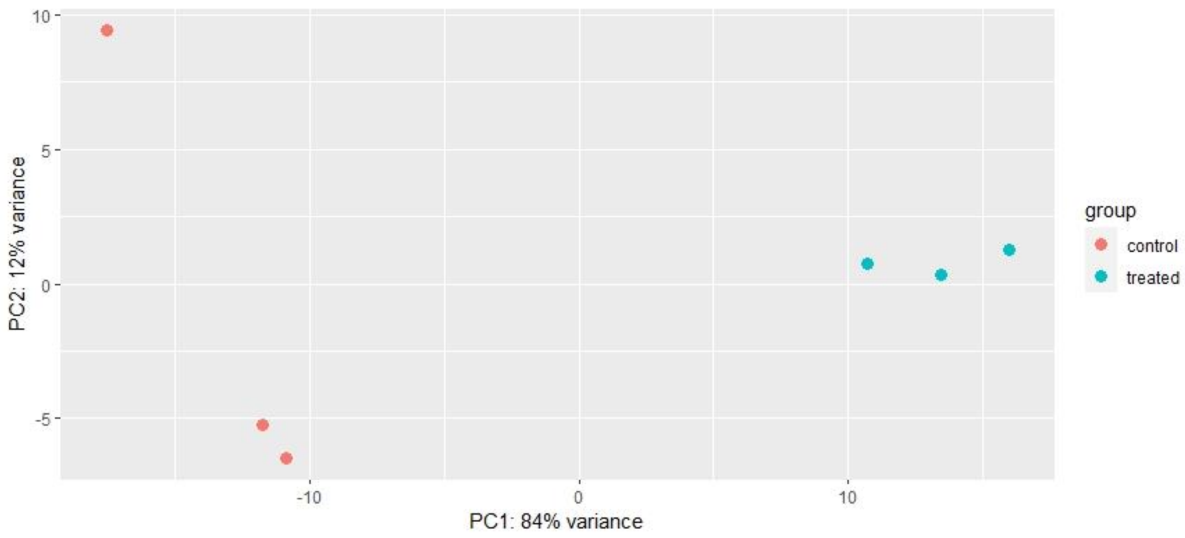
10
5
0
-5
PC2: 12% variance
-10
0
10
PC1: 84% variance
group
control
treated

**Figure S5**

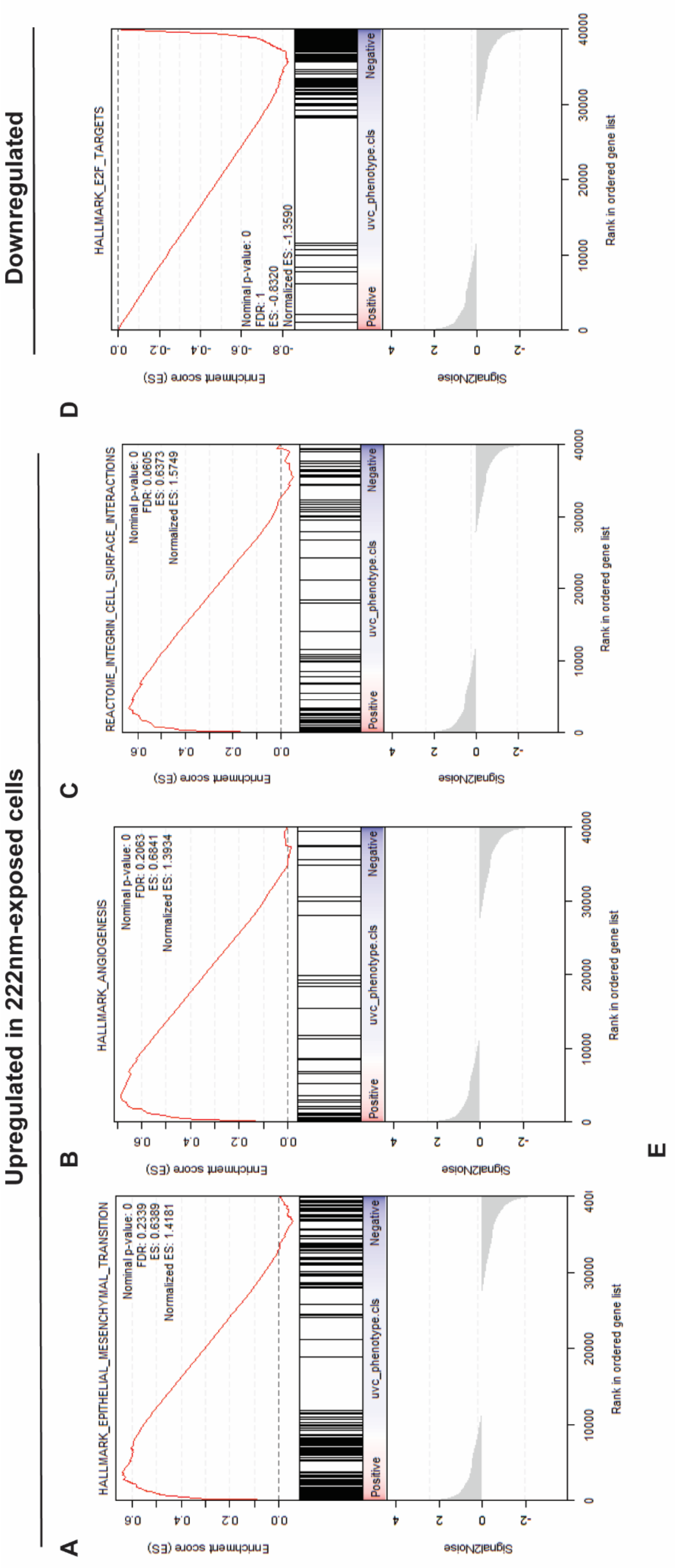
Upregulated in 222nm-exposed cells
Downregulated
A
B
C
D
E
HALLMARK_EPITHELIAL_MESENCHYMAL_TRANSITION
Nominal p-value: 0
FDR: 0.2339
ES: 0.6389
Normalized ES: 1.4181
HALLMARK_ANGIOGENESIS
Nominal p-value: 0
FDR: 0.2063
ES: 0.6841
Normalized ES: 1.3934
REACTOME_INTEGRIN_CELL_SURFACE_INTERACTIONS
Nominal p-value: 0
FDR: 0.0605
ES: 0.6373
Normalized ES: 1.5749
HALLMARK_E2F_TARGETS
Nominal p-value: 0
FDR: 1
ES: -0.8320
Normalized ES: -1.3590
Enrichment score (ES)
Signal2Noise
Positive
uvc_phenotype.cls
Negative
Rank in ordered gene list
